# Cryo-EM Structure of AAA + ATPase Thorase Reveals Novel Helical Filament Formation

**DOI:** 10.1101/2024.11.22.624887

**Authors:** Mohamad Aasif Dar, Robert Louder, Marisol Cortes, Rong Chen, Qianqian Ma, Mayukh Chakrabarti, George K.E. Umanah, Ted M. Dawson, Valina L. Dawson

**Affiliations:** Neuroregeneration and Stem Cell Programs, Institute for Cell Engineering, The Johns Hopkins University School of Medicine, Baltimore, Maryland, United States of America; Department of Neurology, The Johns Hopkins University School of Medicine, Baltimore, Maryland, United States of America; Department of Molecular Biology & Genetics, The Johns Hopkins University School of Medicine, Baltimore, Maryland, United States of America; Department of Biological Chemistry, The Johns Hopkins University School of Medicine, Baltimore, Maryland, United States of America; Department of Biophysics and Biological Chemistry, The Johns Hopkins University School of Medicine, Baltimore, Maryland, United States of America; Solomon H. Snyder Department of Neuroscience, The Johns Hopkins University School of Medicine, Baltimore, Maryland, United States of America; Department of Pharmacology and Molecular Sciences, The Johns Hopkins University School of Medicine, Baltimore, Maryland, United States of America; Department of Physiology, Johns Hopkins University School of Medicine, Baltimore, MD, 21205, USA

## Abstract

The AAA+ (<u>A</u>TPases <u>a</u>ssociated with a variety of cellular <u>a</u>ctivities) ATPase, Thorase, also known as ATAD1, plays multiple roles in synaptic plasticity, mitochondrial quality control and mTOR signaling through disassembling protein complexes like AMPAR and mTORC1 in an ATP-dependent manner. The Oligomerization of Thorase is crucial for its disassembly and remodeling functions. We show that wild-type Thorase forms long helical filaments *in vitro*, dependent on ATP binding but not hydrolysis. We report the Cryogenic Electron Microscopy (cryo-EM) structure of the Thorase filament at a resolution of 4 Å, revealing the dimeric arrangement of the basic repeating unit that is formed through a distinct interface compared to the hexameric MSP1/ATAD1E193Q assembly. Structure-guided mutagenesis confirms the role of critical amino acid residues required for filament formation, oligomerization and disassembly of mTORC1 protein complex. Together, our data reveals a novel filament structure of Thorase and provides critical information that elucidates the mechanism underlying Thorase filament formation and Thorase-mediated disassembly of the mTORC1 complex.

## INTRODUCTION

Thorase is a member of the AAA+ (<u>A</u>TPases <u>a</u>ssociated with a variety of cellular <u>a</u>ctivities) ATPase family. AAA+ ATPases include diverse enzymes that use ATP hydrolysis to drive protein disassembly, nucleic acid remodeling, protein unfolding, protein degradation and other functions(*1–4*). They share a bilobed core of an N-terminal, 200–250 amino acid (aa) nucleotide binding domain, and a C-terminal ~50–80 aa small domain. The nucleotide-binding site has several characteristic signature motifs, including a P-loop, Walker-A and -B, and sensors −1 and −2(*5*). Typically, AAA+ ATPases form a hexameric ring or double-ring structures, wherein the small C-terminal domain is critical for ring formation and substrate engagement; this typically involves threading through the narrow central channel of the ring structure(*6, 7*).

Thorase, also known as ATPase family AAA domain-containing protein 1 (ATAD1), is restricted to multicellular eukaryotes and is expressed in many tissues with the highest expression in the brains and testes(*8*). Thorase uses the energy garnered from ATP hydrolysis to disassemble protein complexes(*8, 9*) with roles in endocytosis and internalization of α-amino-3-hydroxy-5-methyl-4-isoxazolepropionic acid (AMPA) receptors(*8, 10, 11*), maintenance of mitochondrial and peroxisomal integrity(*12–14*), disassembly of the mTORC1 complex(*15*), and has an emerging involvement in human disease(*11, 16, 17*). Thorase plays a critical role in synaptic plasticity by modulating the recycling and trafficking of AMPA receptors on postsynaptic membranes (*8, 11, 14, 16, 17*). Mice with ATAD1 deletions exhibit a seizure-like syndrome caused by an excess of surface-expressed AMPA receptors, resulting in death (*11*). Thorase also plays a critical role in ATP-dependent disassembly of mTORC1 complex, and the disassociation of Thorase from mTOR is critical for reactivating/assembling the mTORC1 complex(*15*). Similarly, experiments with the Walker mutant (E193Q) indicates that it is ATP binding, not the hydrolysis, that is responsible for remodeling and disassembly of the mTORC1 complex (*15*). However, both the precise molecular and structural mechanisms by which Thorase mediates mTORC1 complex disassembly in an ATP-dependent manner, and whether filament formation is vital for this process, is still unknown.

Here, we report that the wild type (WT) ATPase domain of Thorase forms long flexible filaments in the presence of ATP or non-hydrolyzable ATP analogs, and we reveal the cryo-EM structure of the Thorase filament at a resolution of 4 Å. The organization of Thorase subunits within the filament structure represents a unique mode of oligomerization that is novel amongst known AAA + ATPase assemblies, including previously-reported hexameric and dimeric structures of ATAD1-E193Q. Additionally, the crystal structure of ATAD1 yeast homolog, Msp1, was reported not to have a hexamer-like interactions in the crystal packing of wild type MSP1 (*18*). From the Thorase cryo-EM structure, we identify the critical amino acids that are required for filament formation and show that mutation of these residues affect the filament formation and disassembly of mTORC1 complex under *in vitro* conditions. These are essential advances in determining how Thorase directly disassembles protein complexes like mTORC1 and AMPAR in an ATP dependent manner and will provide insight into the function of Thorase/ATAD1.

## RESULTS

### Negative stain analysis of Thorase filament formation

To obtain a homogenous sample for structural analysis, we expressed and purified wild type mouse ATAD1 lacking the first 40 amino acids to exclude the N-terminal transmembrane helix (Fig. 1A). During gel filtration the majority of the protein eluted at the expected volume for the 37 kDa Thorase monomer on Superdex 200 Increase 10/300 column (Fig. 1B). The eluted volume was collected and concentrated before loading on HiLoad 16/600 Superdex 200 column (Fig. S1A) to obtain the highest purity for Thorase monomer, and the peak monomer fraction eluted was used for all experiments herein. SDS-PAGE analysis and western blot of the peak fractions were also carried out (Fig. S1A).

**Fig. 1.**
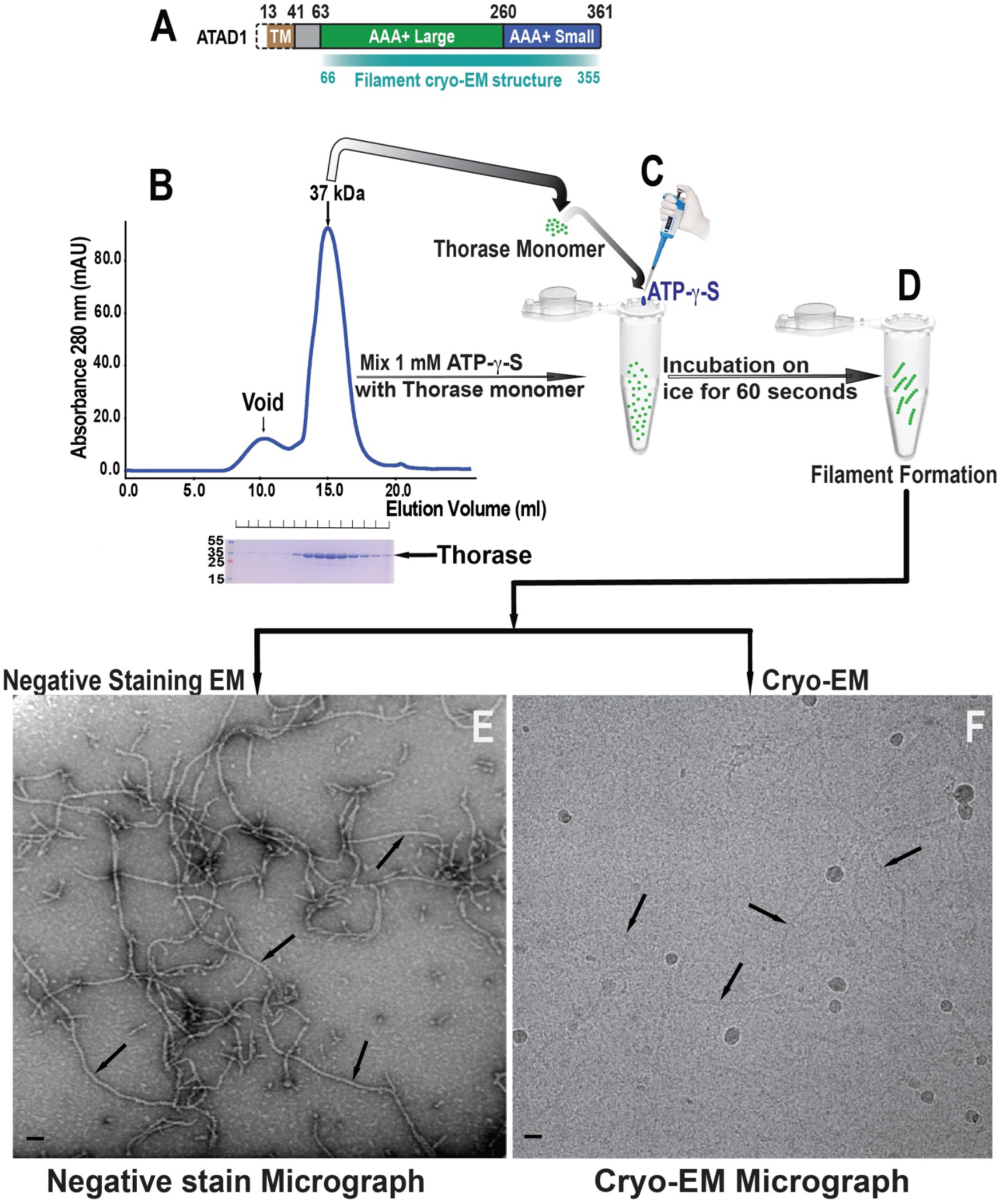
Summary of Purification and Transmission Electron Microscopy of Thorase filament. **(A)** Domain diagram of Mouse Thorase. The construct used in this study comprised residues 41-361. **(B)** Size exclusion chromatography elution profile of Thorase Monomer. **(C and D)** Preparation of Thorase filaments by incubation with non-hydrolysable ATP Analog (ATP-γ-S) **(E)** Negative stain electron micrograph of the Thorase in running buffer with the addition of 1.0 mM ATP-γ-S in which filaments (black arrows) are readily distinguished **(F)** Cryo-EM micrograph of the Thorase in running buffer with the addition of 1.0 mM ATP-γ-S in which filaments in vitreous ice (black arrows) are readily distinguished.

To understand the exact nature of the Thorase oligomeric assemblies, we first performed a negative-stain analysis of the purified protein. The schematic representation for preparation of Thorase filaments for negative staining and cryo-EM experiments are shown in Fig. 1. TEM images of negatively-stained specimens of Thorase prepared subsequent to incubation with 1.0 mM ATP-γ-S for 60 seconds (Fig. 1E) clearly show long filamentous structures with an obvious internal order, with longer ATP-γ-S incubation times resulting in longer filaments (Fig. S1B). ATP and additional ATP analogs (AMP-PNP and ADP-BeF2) were able to support more limited filament formation (Fig. S1C-E).

### Cryo-EM structure of Thorase filament

We next prepared frozen-hydrated cryo-EM specimens of Thorase filaments formed in the presence of ATP-γ-S (Fig. 1F). The resulting cryo-em structure showed Thorase to assemble into a C2-symmetric helical filament with a diameter of ~10 nm, a left-handed twist of 60 degrees, an axial rise of 28.4 Å per subunit, resulting in 6 subunits or ~9 nm rise per helical turn. (Fig. 2A-B; Fig. 3 and Table 1). The top-view suggest a hexameric shape (Fig. 2B – bottom panels). Each layer of the filament is comprised of a Thorase homodimer (Fig. 2C). The micrographs of negatively stained Thorase filaments indicated a relatively large degree of flexibility, and this was further confirmed in the 2D and 3D class averages of the filament segments in cryo-EM (Fig. 3A-B). To reduce the detrimental effect of the flexibility and improve the 3D reconstruction of the complex, we adopted a focused-refinement strategy wherein the boxed filament images were aligned only within a masked region comprising the central 3 layers—corresponding to a single turn of the helical filament— and only C2 point symmetry was applied as opposed to the full helical symmetry (Fig. 3B-C). This strategy allowed a reconstruction of the 3-layer filament segment with resolution range of 4-6 Å. With this reconstruction, we could reliably fit an atomic model for the 6 protomers, encompassing the contiguous range of Thorase residues 66 to 355 (Fig. 2A-B). Surprisingly, this intra-dimer interface is distinct from the subunit interactions observed in the previously reported Thorase hexamer structure as well as the Msp1 dimer structure (Fig. 4). The quality of the cryo-EM map at the dimer interface suffers from the flexibility of the loop comprising Thorase residues 199-204 (Fig. 2D). However, we were able to model the residues in this region using molecular dynamics/flexible fitting (MDFF) after tracing the protein backbone. The resulting model suggests that this interface is largely formed through the electrostatic interactions between an acidic patch formed by E193, D195, D242, and the adjacent ATP-γ-S molecule, with the basic residues R199, R201, and H206 of the opposing protomer (Fig. 2D-E). The point of symmetry lies between R199 residues in the two opposing protomers. Due to high flexibility, the cryo-EM density is weakest for three adjacent serine residues (S202, S203, S204), but their positioning suggests that this part of the loop acts as a lid that interacts with the γ-phosphate of the ATP molecule bound by the opposing protomer (Fig. 2D). The density for bound ATP-γ-S is clearly visible in the cryo-EM maps, and its involvement in forming the dimer interface supports the requirement for ATP binding in filament formation (Fig. 2F). A small interaction between the C-terminal α-helix of one Thorase protomer with the α-helix comprising residues 206-225 of the opposing protomer may also make minor contributions in dimer formation (Fig. 2C).

**Fig. 2.**
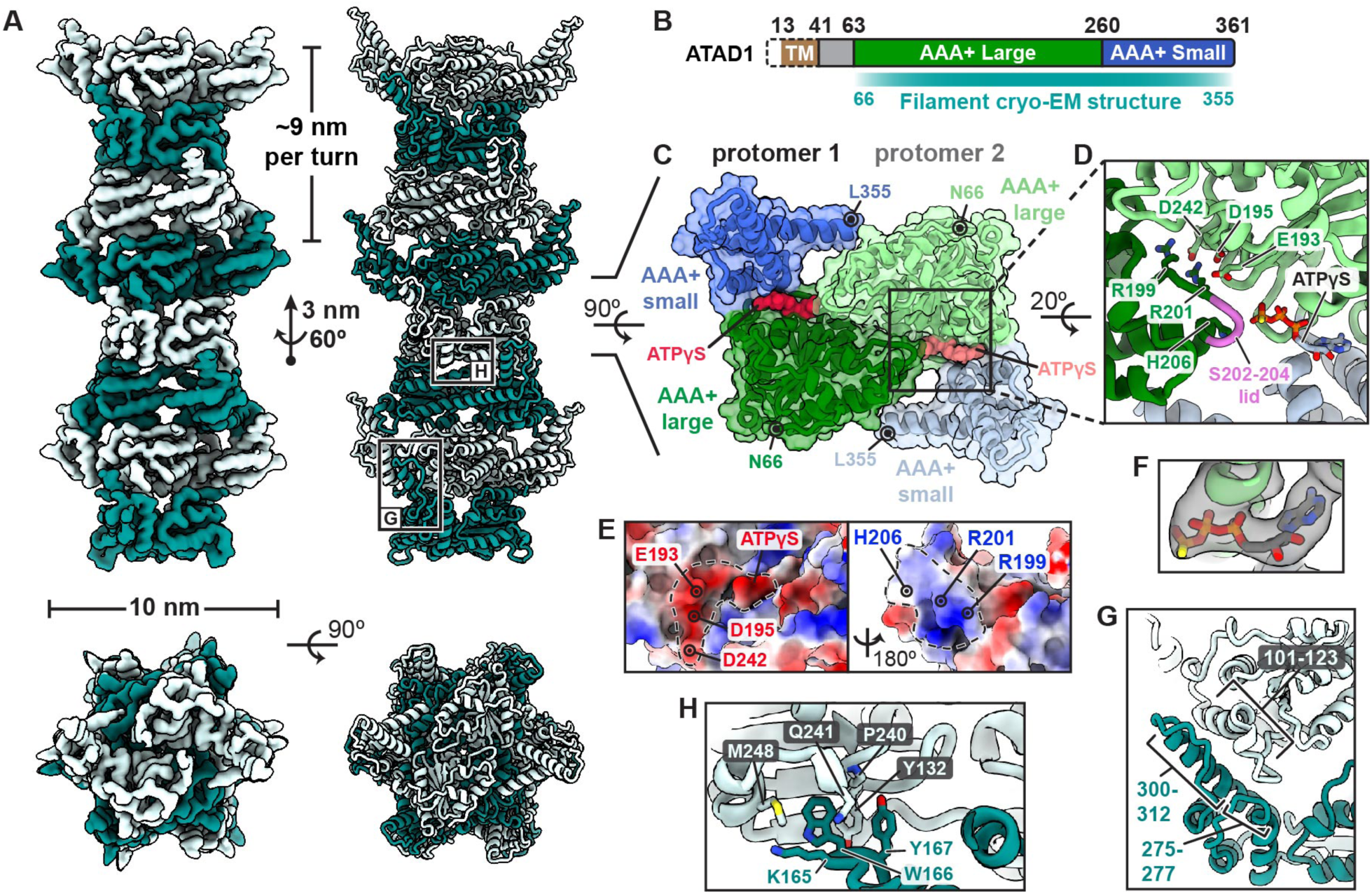
Cryo-EM structure of the Thorase filament. **(A)** Composite 3D reconstruction (left) and model (right) of the Thorase filament generated by fitting and merging multiple copies of the focused-refined map or model, respectively, of a single filament layer into the lower resolution reconstruction of the 8-layer filament segment. Helical parameters are also indicated and Successive layers composed of Thorase dimers alternate between light and dark shades. **(B)** Top and Bottom views of the filament. **(C)** End-on view of a single layer of the Thorase filament illustrating the interface of the dimer that comprises each layer. The large and small AAA+ subdomains are colored in blues and greens, respectively, and ATP-γ-S molecules in reds. The protomer on the right is in lighter shades of the same color scheme. Starting and ending residues for the atomic model are indicated. **(D)** Close up view of the interactions at the primary dimer interface, with participating residues shown in stick representation. Color scheme is similar to panel C. The flexible triple serine loop that forms a lid on top of the bound ATP-γ-S molecule is colored pink. **(E)** Electrostatic potential of the opposing surfaces at the main dimer interface, colored from –10 (red) to +10 (blue) kcal/(mole). **(F)** Cryo-EM density for the bound ATP-γ-S molecule. **(G)** Close-up of longitudinal interactions between filament layers formed at the periphery. **(H)** Close-up of hydrophobic longitudinal interactions located within the filament core. Color scheme in panels F and G match that of panel A.

**Fig. 3.**
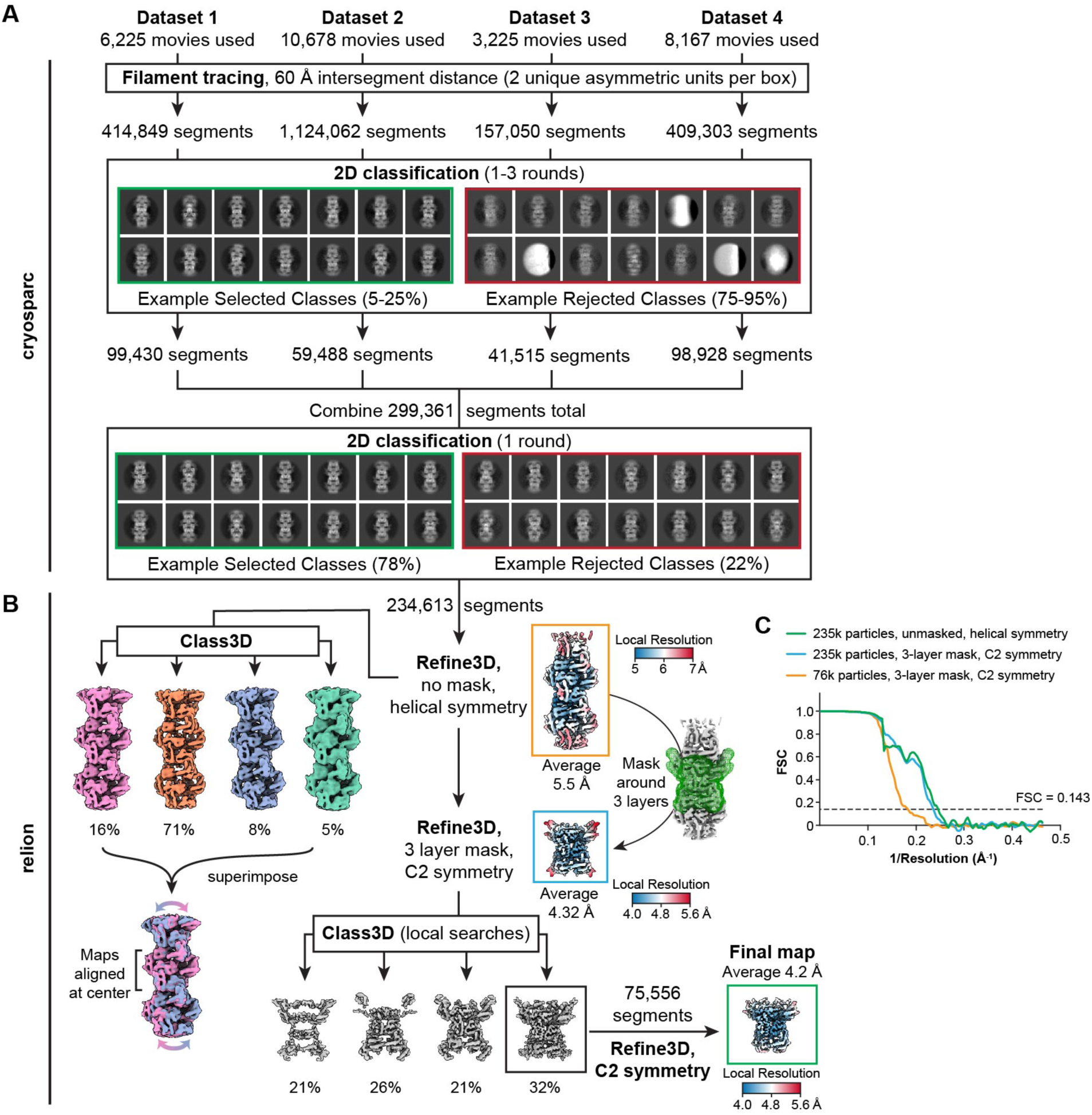
Cryo-EM image processing scheme. **(A)** Four independent data sets were collected and processed independently in CryoSPARC using mainly 2D classification to select for good filament images. A final 2D classification was run on the combined selected filament segment from each dataset. **(B)** Filament segments selected through 2D classification in CryoSPARC were combined and subjected to 3D refinement and classification in Relion. Comparison of 3D averages resulting from classification of unmasked filaments without applied symmetry shows a large degree of conformational heterogeneity within the filament structure (bottom left). Focused refinement within a mask comprising 3 central layers of the filament segments (green) was used to minimize the deleterious effect of filament flexibility and resulted in improved refinement and 3D classification to select for high-resolution filament segments. The final map used for modeling was refined using C2 symmetry and had an average resolution of 4.25 Å. (**C**) Gold standard fourier shell correlation (FSC) plots for the three Refine3D maps, Curve colors correspond to the outline colors for each map.

**Fig. 4.**
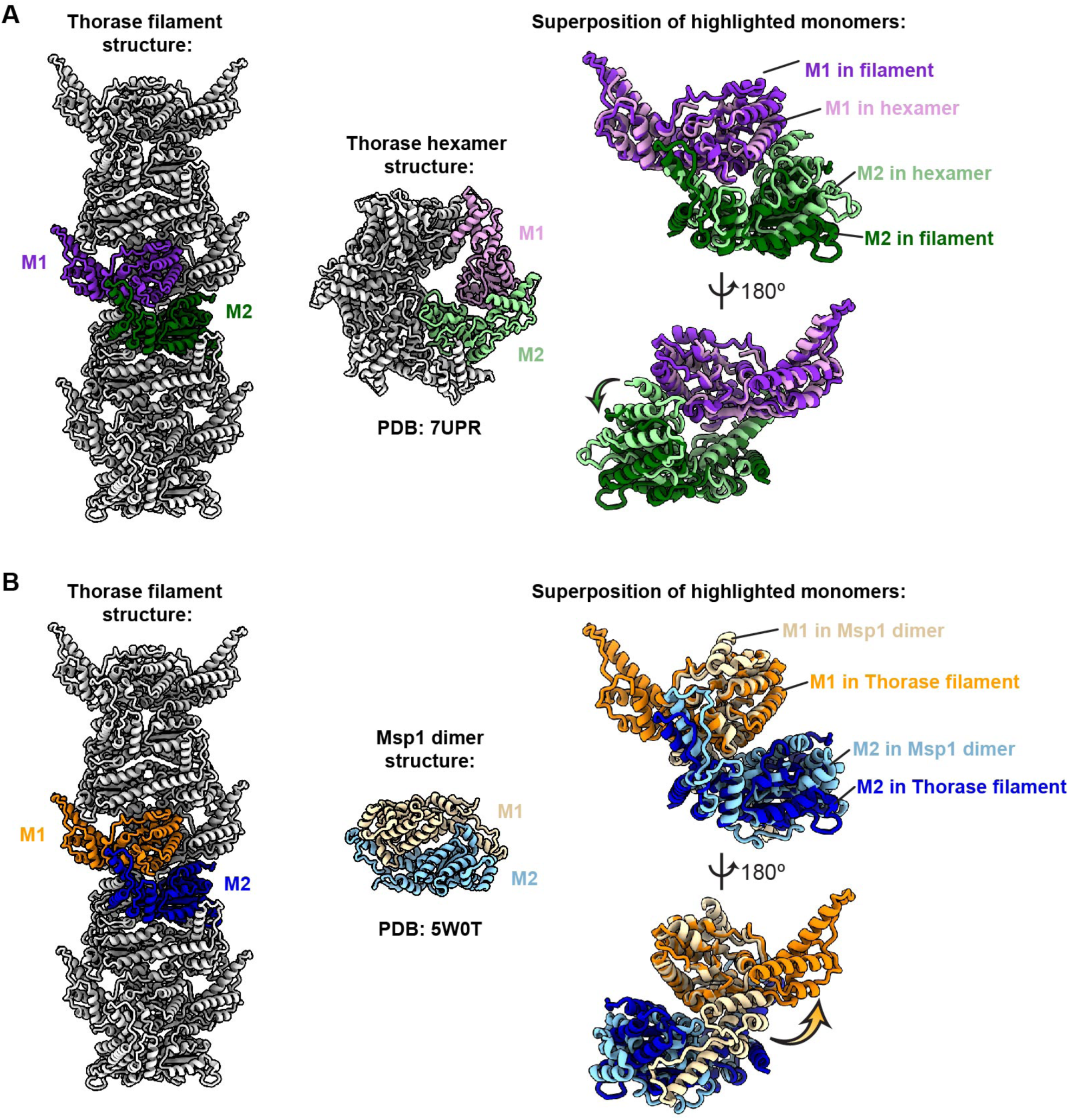
Comparison of the Thorase filament, hexamer, and Msp1 dimer structures. **(A)** Comparison of the longitudinal interface in the Thorase filament with the inter-protomer interface observed in the Thorase hexamer structure. On the right, two protomers from each assembly are shown after superimposing the top protomer, with the colors corresponding to the protomers highlighted in the full filament (left) and hexamer (center) assemblies. **(B)** Comparison of the longitudinal interface in the Thorase filament with the inter-protomer interface observed in the Msp1 dimer structure. On the right, two protomers from each assembly are shown after superimposing the top protomer, with the colors corresponding to the protomers highlighted in the full filament (left) and dimer (center) assemblies.

**Table 1.**
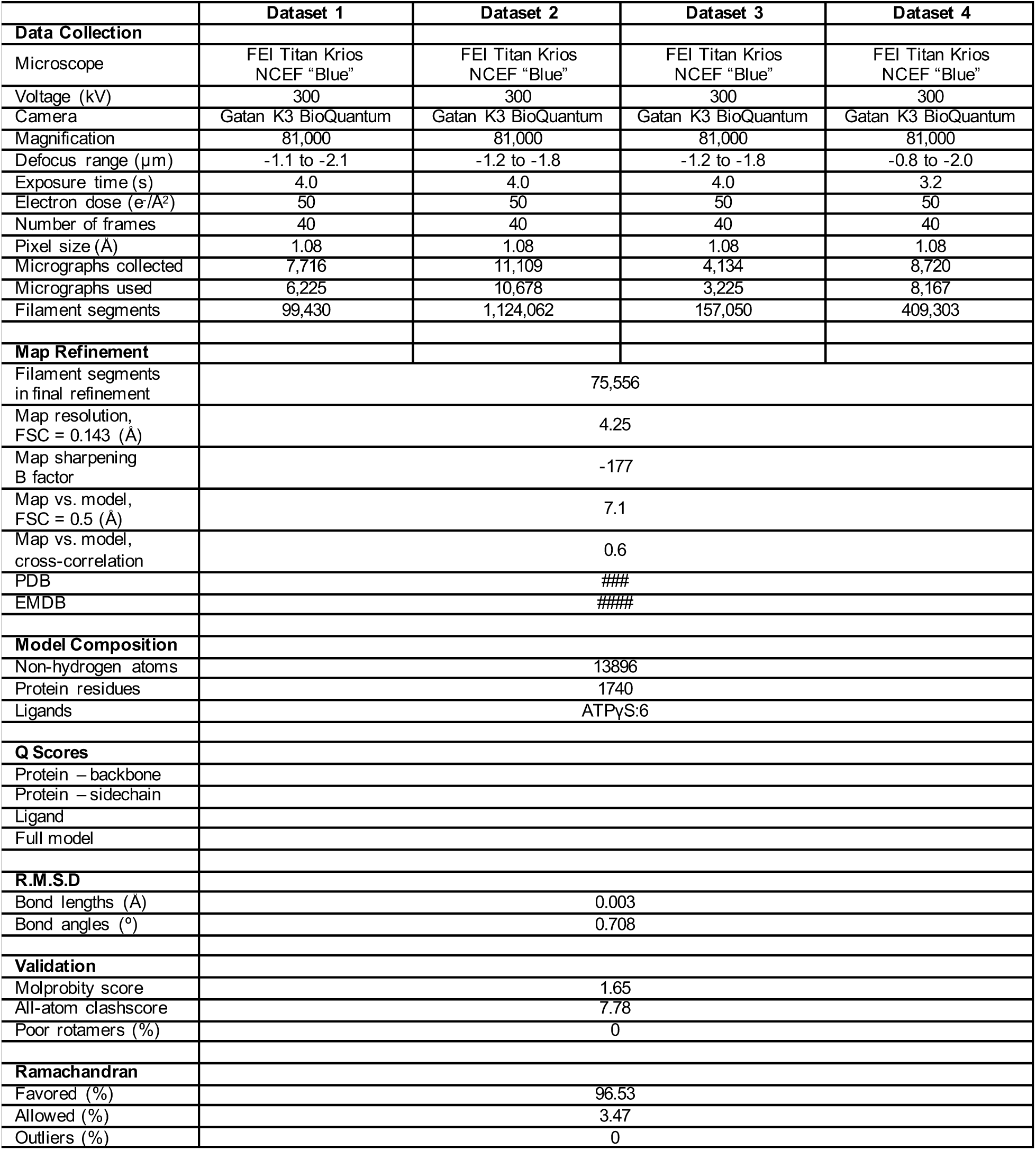
Cryo-EM data collection, refinement and validation statistics.

The filament propagates longitudinally through the stacking of dimers mediated by two interaction regions: longitudinal interactions on the periphery of the filament involve both electrostatic and hydrophobic interactions (Fig. 2G), while those on the interior of the filament involve mainly hydrophobic packing (Fig. 2H). Overall, the residues at the inter-molecular interfaces are highly conserved, suggesting Thorase homologs in other species may also form filaments (Fig. S2).

### Mutagenesis of filament interfaces

Schizophrenia patients from the Ashkenazi Jewish population have a disease-causing mutation that lies in the proximity of these contact sites (E193-D242) that are responsible for filament formation and are highly conserved. Glycine (G222) and glutamate (E290) in this region are likely to be pivotal for the filament formation in an ATP-dependent manner. To prove this point, we mutated Arginine (R) and Aspartic acid (D) to Alanine (A) at contact site 201(R201A), and contact site 195 (D195A). As expected, the mutant proteins did not polymerize, as seen through the negative-stain EM (Fig. 5 F and G). Mutating the conserved residues R304A and M234A in the region also affects filament formation, as observed by negative-stain EM (Fig. 5 D and E). However, R304A forms smaller filaments that are mostly sticking to each other to form smaller aggregates (Fig. 5D).

**Fig. 5.**
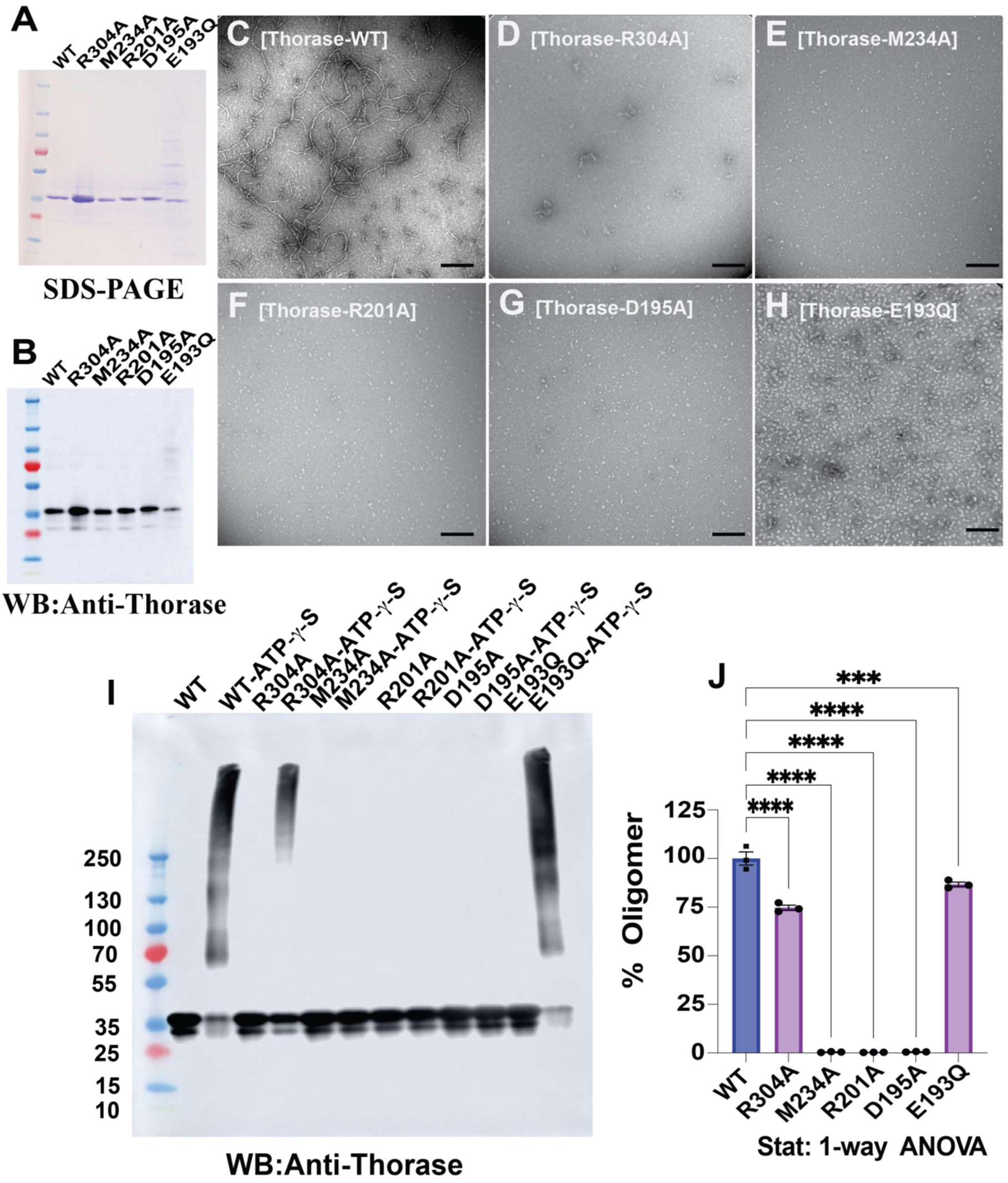
Thorase filament deficient Mutants show defect in ATP-γ-S dependent oligomerization. **(A)** shows SDS-PAGE and **(B)** Western blot of wild type Thorase and Thorase filament deficient mutants respectively. Negative stain electron micrograph of Thorase with 1.0 mM ATP-γ-S **(C)** wild type and Thorase filament deficient mutants: **(D) R304A (E) M234A (F) R201A (G) D195A (H) (I)** Immunoblot shows WT Thorase forms ATP-γ-S dependent oligomer in presence of 1 mM ATP-γ-S. Thorase R304A M234A, R201A and D195A mutants show defects in oligomerization. **(J)** Graphical representation shows quantification of normalized Thorase oligomer in figure I (n=3, experimental repeats). Data = mean ± SEM, One-way ANOVA, ****p <0.0001.

All previously reported structures of Thorase have used the E193Q ATP-trap mutant, which is able to bind but not hydrolyze ATP, thus inactivating the catalytic activity of the enzyme. However, our observation of the involvement of E193 in forming the dimer interface within filamentous wild type Thorase suggests that the mutation of this residue could affect the ability of Thorase to form filaments. To test this, we also purified Thorase E193Q, and we did not observe this mutant forming filaments on negative-stain EM grids (Fig. 5H). Further, we observed that Thorase E193Q elutes as a high molecular weight oligomer through void volume during size exclusion chromatography (Fig. S3A). In the wild type Thorase gel filtration profile, where the majority of the protein eluted at the expected elution volume for the 37 kDa Thorase monomer (Fig. 1B). In addition, it has been already reported by Lan Wang et al that ATAD1-E193Q elutes through the void volume of the Superdex 200 Increase 10/300 column (*19*).

### ATP-γ-S dependent oligomer formation assay

We evaluated oligomer formation for WT Thorase and Thorase filament deficient mutants in the presence of ATP-γ-S using chemical crosslinking agent glutaraldehyde, as previously described (*15, 20, 21*). Thorase WT and filament-deficient mutants R304A, M234A, R201A, D195A and E193Q were purified and SDS-PAGE and western blots are shown in (Fig. 5A and B). Thorase wild type and Thorase filament deficient mutants were cross-linked with glutaraldehyde and immunoblot analyses were also done. The binding of non-hydrolyzable ATP (ATP-γ-S) to WT Thorase facilitates oligomer assembly whereas oligomer formation for Thorase filament-deficient mutants M234A, R201A and D195A were not seen on immunoblots (Fig. 5I). However, R304A forms smaller filaments that are mostly sticking to each other to form aggregates (Fig. 5I). The ATP hydrolysis-deficient Walker B Thorase mutant (E193Q) forms very high molecular weight oligomers (Fig. 5I). In negative staining-EM micrographs, we can see hexameric structures (Fig. S3B), and in immunoblots we can see a band of 250 kDa, corresponding to the molecular weight of the E193Q hexamer, as reported by Lan Wang et al(*19*).

### Thorase filament formation is required for its interaction with mTOR and Raptor

To investigate whether Thorase filament deficient mutants have any effect on binding with mTOR and Raptor, purified recombinant Thorase (WT or mutants) were mixed with whole brain lysates in the presence or absence of ATP or nonhydrolyzable ATP-γ-S to pulldown the mTORC1 proteins (mTOR and Raptor). The schematic representation of recombinant Thorase pulldown of mTORC1 from brain lysate are shown in Fig. 6A. The immunoblots of the purified GST-Tagged Thorase WT and GST-Tagged Thorase filament deficient mutants are shown in Fig. 6B. Although Thorase WT binds both Raptor and mTOR weakly in the absence of nonhydrolyzable Mg^2+^/ATP-γ-S, Thorase WT strongly binds mTOR(Fig. 6C and 6D) in the presence of ATP-γ-S as well as Raptor (Fig. S3C and 3D) in the presence of ATP-γ-S. Interestingly, Thorase filament-deficient mutants in the presence or absence of ATP-γ-S did not show significant changes in binding with Raptor and mTOR (Fig. 6C and 6D, Fig. S3C and 3D).

**Fig. 6.**
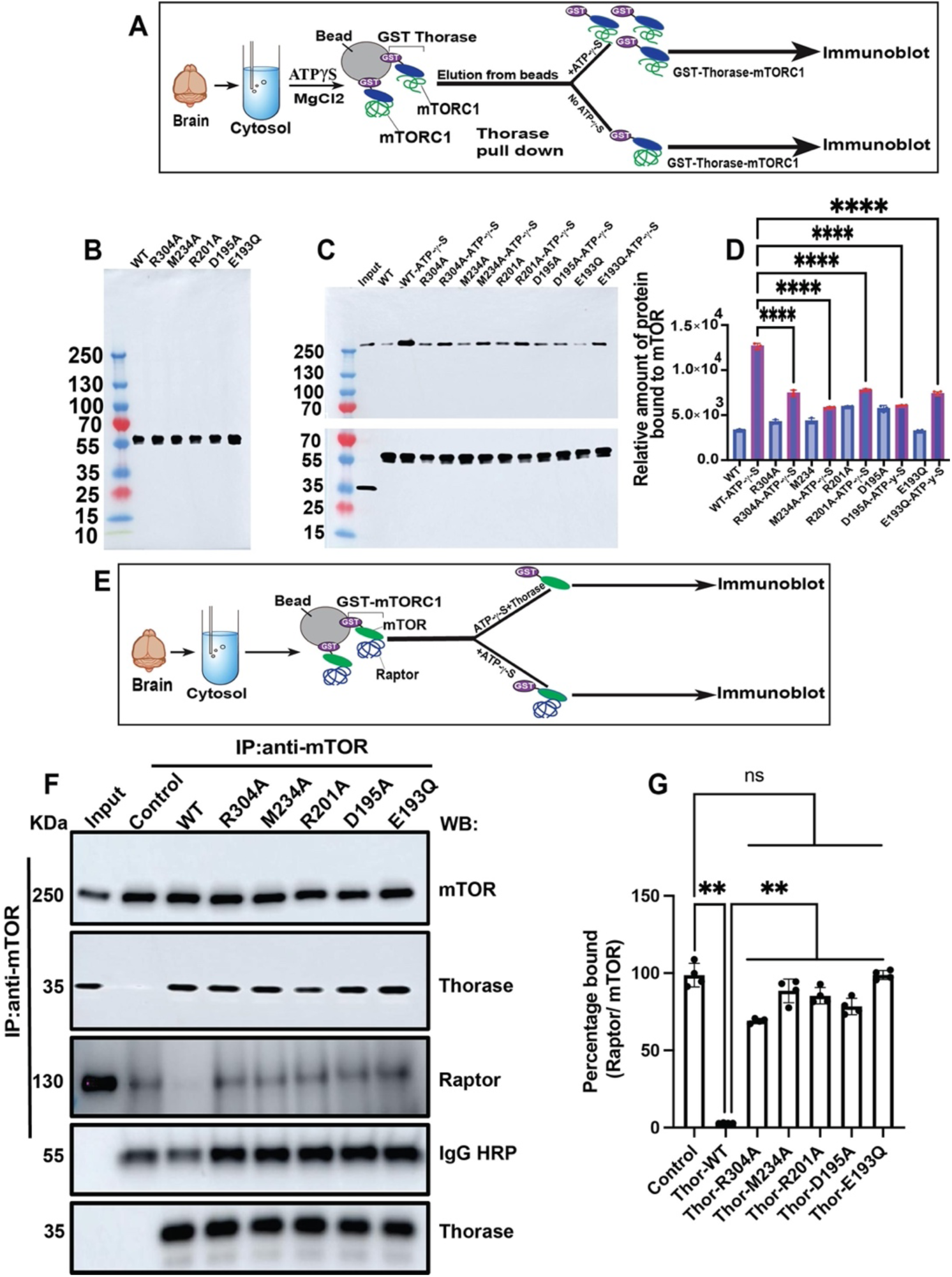
Pull-Down and Disassembly assay from wild type Thorase Brain lysates. **(A)** Schematic representation of recombinant Thorase pulldown from Thorase wild type brain lysate. **(B)** Immunoblot images of purified GST-tagged wild type Thorase and Thorase filament deficient mutants. **(C)** Immunoblot image of immunoprecipitation (IP) of wild type Thorase and Thorase filament deficient mutants in the presence and absence of ATP-γ-S to pulldown mTOR. **D)** Quantification of blots in **C** (n = 3 independent pull-down assays). **(E)** Schematic representation of recombinant Thorase disassembly from Thorase wild type brain lysate. **(F)** Immunoblot images of immunoprecipitation (IP) of mTOR in the presence of wild type Thorase and Thorase Filament deficient mutants **(G)** Graphical representation of quantified blots in A (n = 4 independent IP assays). Data = mean ± SEM, One-way ANOVA, ****p <0.0001.

### Thorase filament formation is required for mTORC1 disassembly

To explore whether Thorase filament-deficient mutants have any effect on mTORC1 disassembly, mTORC1 complexes were prepared from brains of mice. The pulldown mTORC1 on beads was incubated with purified recombinant Thorase (WT or mutants) in the presence or absence of ATP or nonhydrolyzable ATP-γ-S. The schematic representation of disassembly assay from brain lysate is shown in Fig. 6E. The WT Thorase strongly disassembles the mTORC1(Fig. 6 F and G), however Thorase filament-deficient mutants have significantly reduced effect on mTORC1 complex disassembly (Fig. 6 F and G).

## DISCUSSION

This study was inspired by the ambiguity surrounding the oligomeric state of Thorase, which is an important barrier towards revealing the biological role of Thorase. Here, we show that Thorase forms long helical filaments. How rare is this for AAA+ ATPases? While most AAA+ ATPases form hexameric ring structures, Thorase is not the only exception. Prominent, well-studied examples are the MCM2–7 complex, the helicase involved in DNA replication and Torsin A, the AAA+ ATPase involved in the movement of membranes associated with the nuclear envelope and endoplasmic reticulum (*22, 23*). In eukaryotes, the six different subunits of Torsin A form a functional head-to-head double-hexameric ring assembly (*23*). The homologous helicase in archaea of Torsin A, however, only has one distinct subunit and is shown to assemble into oligomers and polymers. It can form helices with a periodicity of 7.2 subunits per turn, and can also form eight-membered rings(*23, 24*). In both assemblies, a wide central channel of ~3–4 nm is observed, which is similar to what has been reported in Torsin A filaments(*25*). However, we do not observe any central channel in Thorase filaments in the cryo-EM structure.

Each layer of the Thorase filament comprises a Thorase dimer, with the intra-dimer interface being distinct from previously observed assemblies of Thorase and its homologs. The filament propagates longitudinally via the addition of dimeric units (layers) on the end, where each subsequent dimer is rotated 30 degrees with respect to the previous one, resulting in 3 layers, or 6 Thorase subunits, per turn. This implies that the Thorase dimer may constitute a stable alternative form of the protein in addition to the previously reported hexameric structure, as the filament is likely to assemble through the addition of dimeric and not hexameric units. While the latter cannot be ruled out, there is little evidence for it in the filament structure.

The high degree of flexibility we observe in the Thorase filament structure can be attributed to the plasticity of the longitudinal protomer interface. Even beyond the filament structure, this interface closely resembles the inter-protomer interface observed within the previously reported hexamer structures of human Thorase and yeast Msp1(*18, 19*). However, a closer distance between the two large AAA+ subdomains in neighboring protomers causes the hexamer to close in on itself in a closed ring instead of propagating linearly (Fig. 4A). The main difference in comparing the same interface in the Msp1 dimer structure (Fig. 4B) can be largely explained by a 180-degree rotation of the small AAA+ subdomain of one of the Msp1 protomers to form the C2 symmetry (Fig. 4B). Thus, it appears that the plasticity of this interface allows Thorase and its homologs to adopt different oligomeric conformations.

Are helical filaments the only self-assembled form of Thorase? If true, one could argue that the interface between adjacent subunits should be distinct from the canonical hexameric interface of other AAA+ ATPases. We compared the Thorase assembly with established hexamers of related AAA+ ATPases: ATAD1E193Q and MSP1E193Q. The difference between the two assemblies is very high, and requires major rotational adjustment of the large versus the small domain.

Protein oligomerization drives many biological processes such as allosteric control of activity, regulation of protein activity by spatial sequestration, control of local concentrations, increased stability against denaturation(*26, 27*) and more. As such, formation of oligomers offers functional, genetic, and physicochemical advantages over monomeric counterparts. A prominent example of this includes oligomeric tripeptidyl peptidase II (TPPII), a cytosolic dimeric enzyme classified as the largest peptidase complex so far(*28*). While TPPII forms a spindle-like rigid structure, the Thorase helix is flexible with no further contacts between neighboring turns. In TPPII, this rigid structure results in restricted access to the active site and the peptidase acts only on smaller and unfolded substrates, showing a 10-fold increased activity after oligomerization. Overall, *in vitro*, no such correlation could be found for oligomeric Thorase, indicating that the functional role of the oligomerization, if any, may only be apparent *in vivo* and could be well explained by cryo-Electron Tomography (cryo-ET).

AAA+ ATPases are known to utilize the energy from ATP hydrolysis to assemble and disassemble protein complexes. To that extent, we previously found that Thorase disassembles GluA2/GRIP1 complexes in the presence of ATP (*8, 11*). In this study we perform traditional immunoprecipitation assays, which indicate that ATP binding, rather than hydrolysis, is required for the Thorase filament formation and interaction between the Thorase filament and mTOR. This interaction disassembles the mTORC1 complex. The structure-guided mutagenesis of Thorase filament deficient mutants shows that the oligomerization and binding of Thorase with mTOR and Raptor is significantly altered. In addition, the oligomerization assay and pull-down assays clearly indicate that filament formation is required for disassembly of the mTORC1 complex by Thorase.

Taken together, we show that Thorase can form filaments in the presence of ATP and ATP analogs, and the structure-guided mutagenesis of Thorase filament-deficient mutants shows that the oligomerization and binding of Thorase with Raptor and mTOR is significantly altered, as revealed in an oligomerization assay and pull-down assays. This shows that filament formation is required for disassembly of the mTORC1 complex by Thorase, even without the ~40 aa hydrophobic N-terminal region present. Going forward, it will be important to further dissect the ATP hydrolysis cycle and to study the protein *in situ* at higher resolution as well as elucidate the structure of the Raptor/mTOR-Thorase filament complex structure by cryo-EM. Finally, it would be very exciting to resolve the structure of Thorase filaments with N-terminal 40 residues inside the cell by labelling the Thorase with a fluorescent tag for expression in HEK cells, followed by vitrification and FIB milling. This would permit an *in situ* structure to be determined by Cryo-ET.

### Limitations of the study

One important, potential limitation of our study is that we used an N-terminally truncated Thorase construct in our structural analysis making it possible that the structures are fully representative of full-length Thorase. The most direct test would be to do the same study with the full-length Thorase. The missing 40 N-terminal residues (aa 1–40) are predicted to be solvent exposed in the filament structure, and thus, they should not block assembly. This N-terminal peptide is hydrophobic and is suggested to interact with membranes mediating AMPAR trafficking (*8*). Unfortunately, the full-length protein is ill-behaved in our hands and it aggregates during purification because of the hydrophobic N-terminal transmembrane domain, thus preventing us from doing meaningful experiments. The main function of N-terminal peptide is an interesting subject to explore in future studies by using Cryo-ET.

## Acknowledgments

We are grateful to Duncan Souza for training and guidance for screening cryo-EM grids at Johns Hopkins Medicine.

## Funding

This research was supported by NIH/NIDA DA000266, DA044123 to T.M.D. and V.L.D and NIH/NINDS and NS099362 to GKEU. TMD is the Leonard and Madlyn Abramson Professor in Neurodegenerative Diseases. This research was, in part, supported by the National Cancer Institute’s National cryo-EM Facility at the Frederick National Laboratory for Cancer Research under contract 75N91019D00024.

## Author contributions

Conceptualization: MAD, GKEU, TMD, VLD; Methodology: MAD, RL, MC, RC, QM, MC; Investigation: MAD, RL, TMD, VLD; Visualization: RL, MAD; Supervision: TMD, GKEU, VLD; Writing – Original draft: MAD, RL; Writing – review and editing: GKEU, TMD, VLD

## Competing interests

The authors declare no competing interests.

## MATERIALS AND METHODS

### Protein expression and purification

Mouse Thorase (residues 41–361), N-terminally fused with a human rhinovirus 3C protease cleavable GST tag, was cloned into an ampicillin resistant pGEX-6P-1 vector (addgene). The E. coli strain BL21(DE3) (Thermo Fischer Scientific) was transformed with the GST-Thorase construct. Cells were grown at 37°C in Luria-Bertani (LB) broth medium supplemented with 100 µg ml^-1^ ampicillin, until an optical density (OD_600_) of 0.6–0.8 was reached, shifted to 4 °C for 1 hour and induced overnight at 16 °C with 0.2 mM isopropyl β-D-1-Thiogalactopyranoside (IPTG). The bacterial cultures were harvested by centrifugation, resuspended in lysis buffer (100 mM Tris HCl pH 7.5, 200 mM NaCl, 5.0mM MgCl2 and 5% glycerol) and lysed at 15000 psi with a high-pressure homogenizer (LM20 Microfluidizer, Microfluidics) The lysate was immediately mixed with 0.1 M phenylmethanesulfonyl fluoride (PMSF) (50 µl per 10 ml lysate) and 1X of Protease Inhibitor Cocktail (Thermo Fischer Scientific), and cleared by centrifugation at 12000 RPM for 30 minutes. The soluble fraction was gently mixed with Pierce™ Glutathione Agarose resin (Thermo Fischer Scientific) for 3 hours at 4°C on a rotating mixer (Benchmark). After washing with GST-binding buffer (100 mM Tris-Cl pH 7.5, 200 mM NaCl, 5.0 mM MgCl2, 5.0 mM TCEP and 5% glycerol) at least three times, the PreScission buffer containing PreScission Protease (General Electric) was added to the Glutathione agarose resin and incubated for 4.0 hours at 4°C on a Bench rocker (Benchmark). After incubation the mixture was centrifuged at 1000 RPM for 5 minutes at 4°C. The supernatant was collected and concentrated approximately 3-fold using a 10 kDa cut-off Amicon Ultra-15 centrifugal filter unit concentrator (Millipore). Approximately 2.0 mL of concentrated protein was subjected to size-exclusion chromatography using the Äkta Pure system and a Superdex 200 increase 10/300 gl column pre- equilibrated with running buffer (100 mM Tris-Cl pH 7.5, 200 mM NaCl, 5.0 mM MgCl2 and 5.0 mM TCEP). Peak fractions were collected and analyzed for purity and composition with SDS-PAGE and the fractions containing the Thorase with purity > 70% were again concentrated using Amicon Ultra-15 centrifugal filter unit (10-kDa molecular mass cut-off; Merck, Darmstadt, Germany). Approximately 2.0 mL of concentrated protein was subjected to size-exclusion chromatography using the Äkta Pure system and a HiLoad 16/600 Superdex 200 gl column pre-equilibrated with running buffer. The eluate was collected in fractions and analyzed for purity and composition with SDS-PAGE and the purity of the protein was >95%. These fractions containing pure protein were pooled and concentrated using an Amicon Ultra-15 centrifugal filter unit (10-kDa molecular mass cut-off; Merck, Darmstadt, Germany) until a total protein concentration of 1.20 mg/mL was reached. After concentration, the protein was stored in 20 µl of aliquots in 1.5 ml Protein LoBind Tubes (Eppendorf) and was flash-frozen in liquid nitrogen for future use.

### Negative stain Electron Microscopy

All the reaction steps were carried out at 4 °C. To prepare Thorase filaments 1.0 µl of fresh 0.1 M ATP-γ-S (Millipore Sigma) dissolved in running buffer was added to 20 µl of protein having concentration of 0.15 mg/ml for Negative staining EM and 1.20 mg/ml for cryo-EM, and the mixture was gently mixed by pipette and incubated on ice for 60 seconds. After incubation filaments were ready for both negative staining as well as for cryo-EM experiments. Carbon Square Mesh, Cu, 400 Mesh grids, (Electron Microscopy Sciences) were glow discharged for 45 s before 5.0 µL of Thorase filaments (A280 = 0.15) were applied. Samples were left on the grids for 1 min, blotted, and then stained with 2% uranyl acetate for 60 s, blotted, and air dried. The grids were imaged on a Hitachi 7600 TEM, 80 keV microscope at the Johns Hopkins University, School of Medicine EM facility.

### Cryo-Electron Microscopy

To prepare the cryo-EM specimen, 4 µl of the reaction mixture containing Thorase filaments was placed on a glow-discharged Lacey Carbon (300 mesh cu) and flash-plunged into liquid ethane cooled at liquid-nitrogen temperature with a FEI Vitrobot Mark IV with a blotting time of 4.0 s and a blotting force of 3 in an environment with 100% humidity and a 4 °C temperature. The grids were transferred to a Titan Krios electron microscope (FEI) operated at an acceleration voltage of 300 kV and equipped with a K3 direct electron-counting camera (Gatan). The grids were screened and the best grids were selected for data collection. The screened grids were sent to The National Cancer Institute (NCI) in Bethesda, Maryland, USA for data collection (Table 1). Movies of the specimens were recorded manually at a nominal magnification of 81,000 with K3 Nanoprobe EFTEM mode, thus yielding a pixel size of 1.08 Å, with the NCI-Image4 data-collection user interface(*29*). The specimen was exposed to a dose rate of approximately 12.50 e-/Å2/sec. pixel and total exposure time of 4.0 s, with total of 40 frames per exposure with a defocus value ranging from −0.75 to −2.0 µm.

### Cryo-EM image processing

Around 28,295 movies collected across four different sessions were combined for the final cryo-EM structure, after eliminating movies without picked filaments. Movies from each dataset were initially processed independently in CryoSPARC (*30*), starting with patch motion correction and patch CTF estimation. Motion-corrected micrographs were then used for template-free filament tracing in CryoSPARC, selecting for traces longer than 48 nm. Filament segments were extracted in 276.5 Å (256 pixel) boxes with a separation distance of 60 Å between segments, corresponding to roughly 10 filament layers total and 2 unique layers (4 Thorase subunits) per boxed segment. Filament segments were subjected to 2D classification in CryoSPARC, and three well-defined filament class averages representing different filament views were used for template-based filament tracing. Filament segments from template tracing were again extracted using the same parameters, subjected to several rounds of 2D classification followed by selection of particles belonging to well-defined class averages.

The helical parameters for the filament were first approximated empirically and were further refined using three-dimensional (3D) helical refinement with symmetry searches turned on. The 3D helical refinement from the higher quality segments resulted in reconstructions with similar resolutions for each data set. Thus, we combined the filament segments from all data sets and sorted them together by another round of 2D classification, and particles from class averages with obvious kinks in the filament structure were excluded from further processing. A total of 234,613 boxed filament segments were then used for 3D classification and refinement in Relion (*31*). Conversion of the data files from CryoSPARC to Relion format was done using pyem (*32*). Filament segments were first aligned globally in Relion by 3D refinement with helical symmetry applied. Initial 3D classification of the aligned filament segments without applied symmetry resulted in 3D class volumes that exhibited a large degree of conformational flexibility within the filament structure (Fig. 3). To reduce the detrimental effect of this flexibility on the resolution of the 3D reconstruction, we refined the segment alignments in a focused region that included only the central 3 layers of the filament (6 Thorase subunits). This focused refinement resulted in a reconstruction of markedly improved resolution and map quality. Further improvement was achieved through 3D classification of the segments in the focused region with local translational and angular searches, selection of the highest-resolution class, and another round of focused refinement of the higher resolution segments. The final focused-refined map resolution ranged from 4-6 Å with an average of 4.2 Å.

Map sharpening was performed with deepEMhancer (*33*). Various amplitude corrected maps were employed for model building. Conversion between cryoSPARC and RELION were performed with pyem.

### Model building and refinement

The initial model for human Thorase was obtained from the AlphaFold Protein Structure Database (AF-Q8NBU5-F1_v4) and low confidence regions were trimmed (pLDDT < 50) prior to rigid body fitting 6 copies into the 3-layer focused-refined cryo-EM density with UCSF Chimera (v1.13)(*34*). Subsequently, the model was flexibly fit and interactively refined using ISOLDE (*35*) within Chimera X (v1.3) (*36, 37*) and Coot(*38*). All models were subject to real space refinement and validation in PHENIX (*39*) and MolProbity (*40*).

### ATPase Activity Assay

The ATP hydrolysis activity was determined by using the ADP Colorimetric Assay Kit (Millipore Sigma) following manufacturer’s instruction with some modifications. Briefly, the recombinant proteins 20 ug in a 50μl system were incubated with 200 μM ATP for 30 min at 37°C for ATP hydrolysis to ADP. To check ATP degradation to ADP that were not due to enzymatic hydrolyses the ATPase buffer (without protein) containing 200 μM ATP and Thorase E193Q containing 200 μM ATP were included as control. The samples were incubated with ADP reaction mix buffer for 30 min at room temperature in the dark and the absorbances at 590 nm for each sample (in triplicates) were measured to determine the amount of ADP present in the samples. The rate of ATP hydrolysis was determined by the rate of ADP formation.

### Oligomer formation assays

To evaluate the oligomer formation of Thorase and Thorase filament deficient mutants, purified recombinant proteins (5 μg in 100 μl) were treated with 1 mM ATP-γ-S in modified HEN buffer (250 mM HEPES, 2 mM MgCl2, 0.1 mM Neocuproine, 0.5% Nondiet P-40) at 4°C. Oligomer formation upon ATP-γ-S binding was examined by chemical cross-linking with 0.02% glutaraldehyde at room temperature for 5 minutes as previously described (Babst et al., 1998). The cross-linked products were TCA-precipitated, washed with acetone and analyzed by SDS– PAGE and Immunoblotted with anti-Thorase antibody.

### Pull Down Assay

For pull down of mTOR and Raptor, recombinant GST-Thorase (wild type and filament deficient mutants) were purified on GST-beads and mixed with mouse whole brain lysates in the presence of 2mM ADP, ATP, or ATP-γ-S in buffer A and incubated with mixing for 2 h (*41*). The beads were then extensively washed with buffer A containing 1mM ADP, ATP, or ATP-γ-S. The beads were resuspended in 2X SDS-PAGE Laemmli buffer and eluted samples were resolved on 10% SDS-PAGE. Immunoblotting was used to confirm the presence of mTOR and Raptor proteins bound to GST tagged wild type Thorase and Thorase filament deficient mutants.

### Disassembly Assay

Thorase disassembly of the mTORC1 complex was examined by immunoprecipitating Raptor and mTOR from isolated mouse brain lysate as previously described with some modification (Umanah et al., 2017; Zhang et al., 2011). Briefly, freshly mouse brains were homogenized in binding buffer (25 mM HEPES-KOH (pH 7.8), 150 mM NaCl, 3 mM MgCl2, 1 mM DTT) containing protease inhibitors, and Triton X-100 was added to a final concentration of 1% and incubated at 4°C (with mixing by end-to-end rotation) for 1 hour. Samples were centrifuged at 15,000 x g for 30 min and supernatant was collected. The supernatant, recombinant wild type Thorase and Thorase filament-deficient mutants were incubated with Protein G Magnetic Beads (Invitrogen) pre-bound with anti-mTOR antibodies plus 2 mM ADP or ATP or ATP-γ-S at 4°C (with mixing by end-to-end rotation) for 1 hour. The samples were then transferred to room temperature and incubated (with mixing by end-to-end rotation) for 1 hour. Beads were washed 3 times with binding buffer (with mixing by end-to-end rotation) for 2 minutes per wash. After the last wash bound proteins were eluted from beads using 2x SDS-PAGE Laemmli buffer with DTT. The eluted proteins were resolved on 10% SDS-PAGE and immunoblotted using anti-Raptor, anti-mTOR and anti-Thorase antibodies.

### Data and materials accessibility

The helical EM reconstruction of the Thorase filament is available from the Electron Microscopy Data Bank under accession code EMD-43640. The focused-refined 3-layer C2-symmetric EM map and corresponding atomic coordinates have been deposited to the Electron Microscopy Data Bank and Protein Data Bank under accession codes EMD-43639 and PDB ID 8VXT, respectively. All data are available in the main text or the supplementary materials. Source data files and construct materials related to this work are available upon request.

---

**Extended Data Fig. 1.**
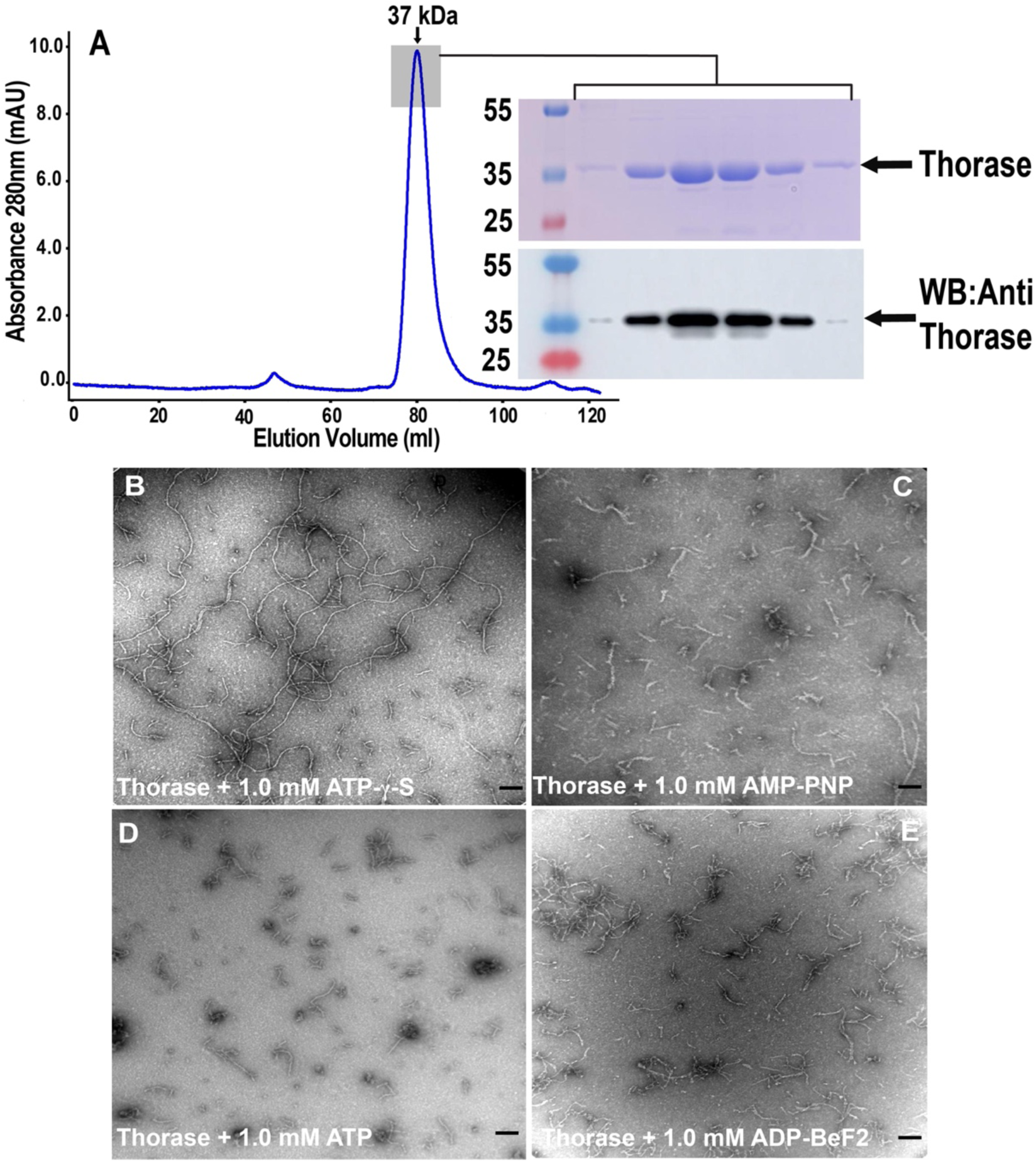
Purification and biochemical characterization of Thorase filaments. **(A)** Size exclusion chromatography elution profile of Thorase. The experiment was performed on a HiLoad 16/600 Superdex 200 column in running buffer containing 100 mM Tris HCl pH 7.5, 150 mM NaCl, 5 mM MgCl_2_ and 5 mM TCEP. SDS-PAGE analysis of the fractions is shown on right side of the chromatogram as well as western blot of the fractions are also shown. **(B)** Negative stain electron micrograph of the Thorase in running buffer with the addition of 1.0 mM ATP-7-S. **(C)** Negative stain electron micrograph of the Thorase in running buffer with the addition of 1.0 mM AMP-PNP. **(D)** Negative stain electron micrograph of the Thorase in running buffer with the addition of 1.0 mM ATP. **(E)** Negative stain electron micrograph of the Thorase in running buffer with the addition of 1.0 mM ADP-BeF2. Scale bar, 200 nm.

**Extended Data Fig. 2.**
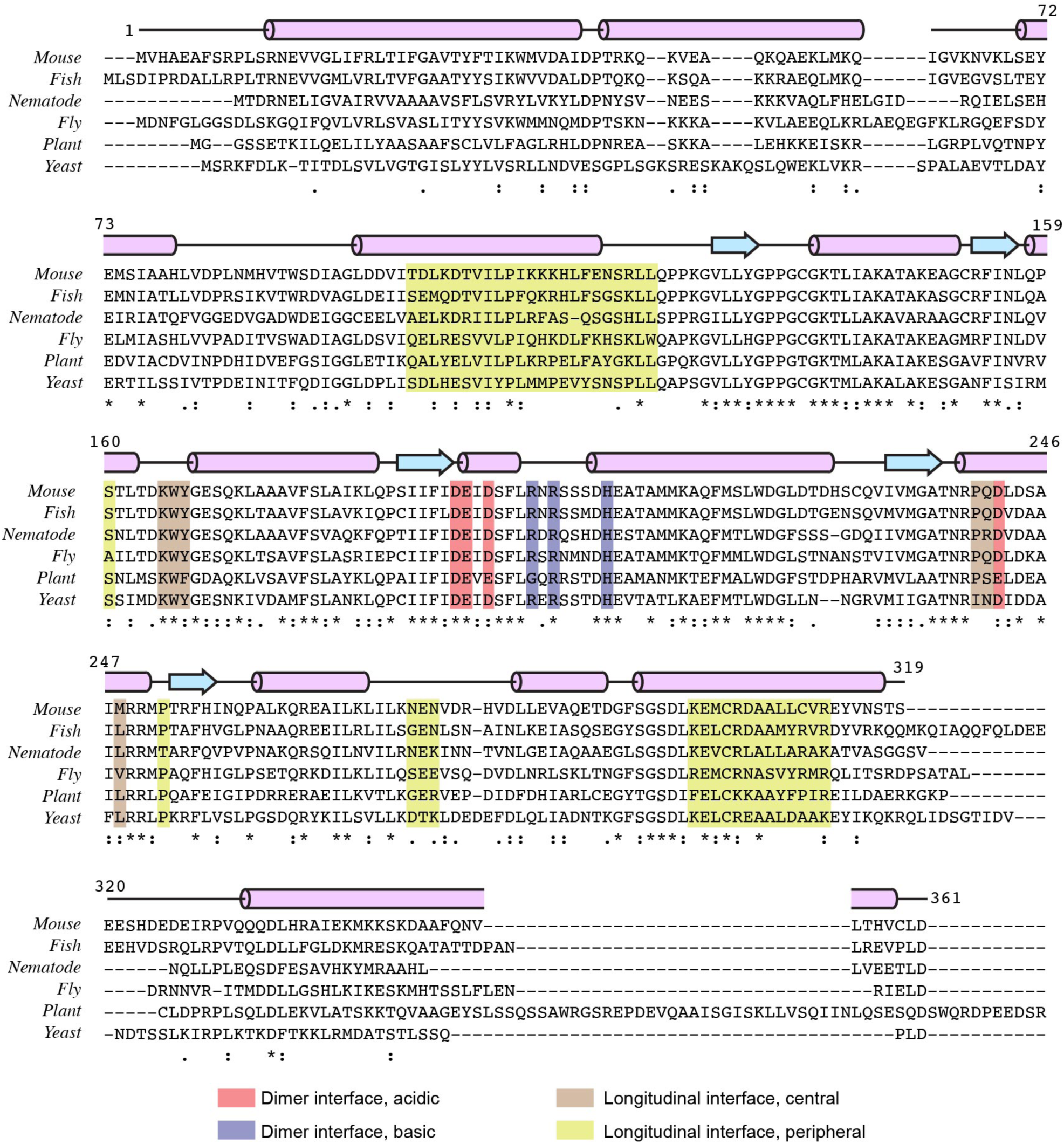
Multiple sequence alignment of Thorase and its homologs in other model eukaryotes. Regions of inter-protomer contacts observed within the filament structure are highlighted with different colors, according to the legend at the bottom. Hs, Homo sapiens; Dr, Danio rerio; Ce, Ceanorhabditis elegans; Dm, Drosophila melanogaster; At, Arabidopsis thaliana; Sc, Saccharomyces cerevisiae.

**Extended Data Fig. 3.**
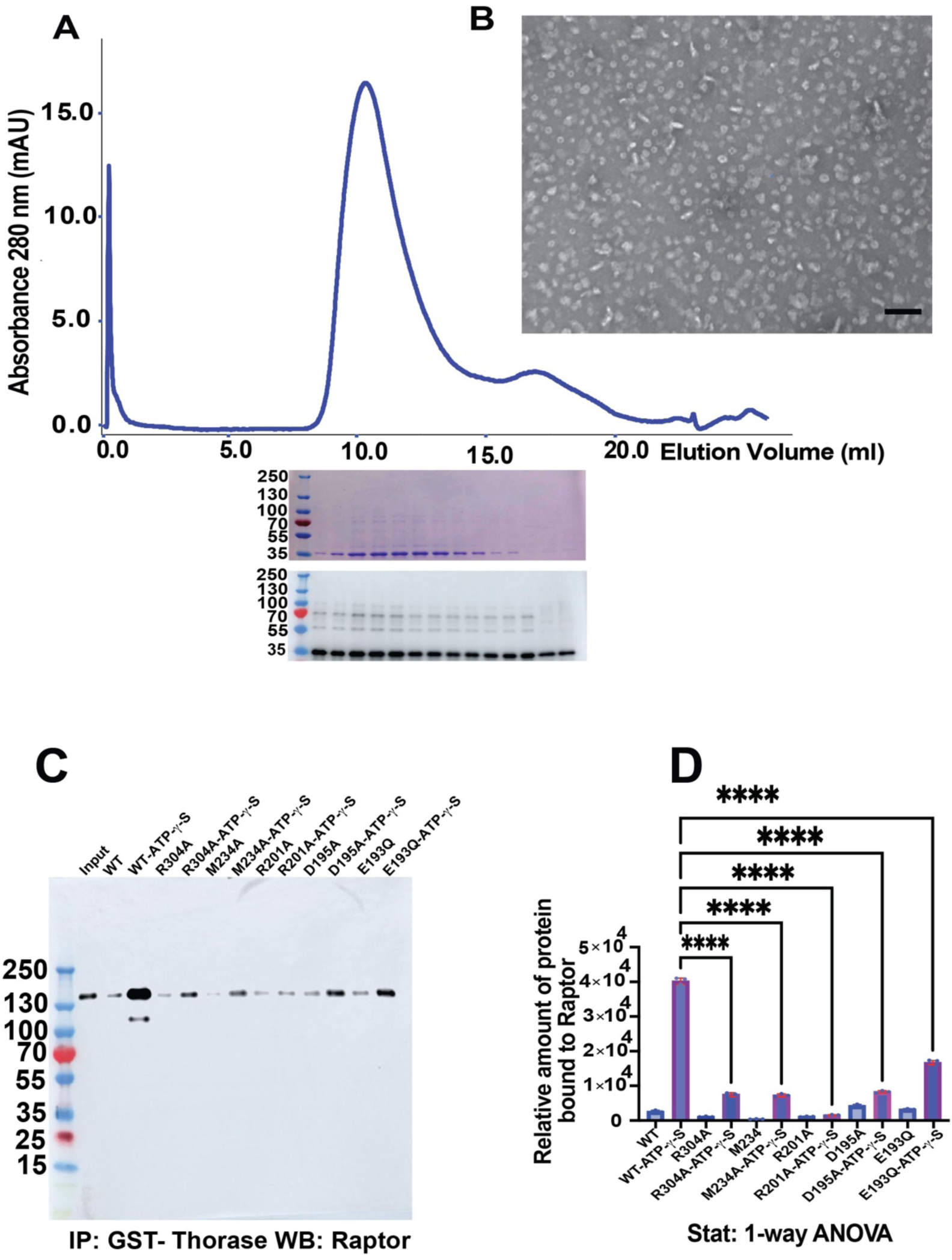
Purification and negative staining of Thorase ATP trap mutant E193Q. **(A)** Size exclusion chromatography elution profile of Thorase ATP Trap mutant E193Q. The experiment was performed on a Superdex 200 Increase 10/300 Column in running buffer containing 100 mM Tris HCl pH 7.5, 150 mM NaCl, 5 mM MgCl_2_ and 5 mM TCEP. SDS-PAGE and Western blot analysis of the fractions of the two peaks collected is shown below the chromatogram. **(B)** Negative stain electron micrograph of the Thorase ATP Trap mutant E193Q in running buffer with the addition of 1.0 mM ATP-γ-S. Scale bar, 100 nm **(C)** Immunoblot image of immunoprecipitation (IP) of wild type Thorase and Thorase filament deficient mutants in the presence and absence of ATP-γ-S to pulldown Raptor. **(D)** Quantification of blots in B (n = 3 independent pull-down assays).

## Notes

### Competing Interest Statement

The authors have declared no competing interest.

## REFERENCES

1. J. P. Erzberger, J. M. Berger, Evolutionary relationships and structural mechanisms of AAA+ proteins. Annu Rev Biophys Biomol Struct 35, 93–114 (2006).

2. L. M. Iyer, D. D. Leipe, E. V. Koonin, L. Aravind, Evolutionary history and higher order classification of AAA+ ATPases. J Struct Biol 146, 11–31 (2004).

3. T. A. Sysoeva, Assessing heterogeneity in oligomeric AAA+ machines. Cell Mol Life Sci 74, 1001–1018 (2017).

4. M. Jessop, J. Felix, I. Gutsche, AAA+ ATPases: structural insertions under the magnifying glass. Curr Opin Struct Biol 66, 119–128 (2021).

5. P. Wendler, S. Ciniawsky, M. Kock, S. Kube, Structure and function of the AAA+ nucleotide binding pocket. Biochim Biophys Acta 1823, 2–14 (2012).

6. K. Nyquist, A. Martin, Marching to the beat of the ring: polypeptide translocation by AAA+ proteases. Trends Biochem Sci 39, 53–60 (2014).

7. A. O. Olivares, T. A. Baker, R. T. Sauer, Mechanistic insights into bacterial AAA+ proteases and protein-remodelling machines. Nat Rev Microbiol 14, 33–44 (2016).

8. J. Zhang et al., The AAA+ ATPase Thorase regulates AMPA receptor-dependent synaptic plasticity and behavior. Cell 145, 284–299 (2011).

9. C. Dai et al., Functional identification of neuroprotective molecules. PLoS One 5, e15008 (2010).

10. M. Pignatelli et al., Synaptic Plasticity onto Dopamine Neurons Shapes Fear Learning. Neuron 93, 425–440 (2017).

11. G. K. E. Umanah et al., Thorase variants are associated with defects in glutamatergic neurotransmission that can be rescued by Perampanel. Sci Transl Med 9, (2017).

12. Y. C. Chen et al., Msp1/ATAD1 maintains mitochondrial function by facilitating the degradation of mislocalized tail-anchored proteins. EMBO J 33, 1548–1564 (2014).

13. V. Okreglak, P. Walter, The conserved AAA-ATPase Msp1 confers organelle specificity to tail-anchored proteins. Proc Natl Acad Sci U S A 111, 8019–8024 (2014).

14. J. Prendergast et al., Ganglioside regulation of AMPA receptor trafficking. J Neurosci 34, 13246–13258 (2014).

15. G. K. E. Umanah et al., AAA + ATPase Thorase inhibits mTOR signaling through the disassembly of the mTOR complex 1. Nat Commun 13, 4836 (2022).

16. R. C. Ahrens-Nicklas et al., Precision therapy for a new disorder of AMPA receptor recycling due to mutations in. Neurol Genet 3, e130 (2017).

17. J. Piard et al., A homozygous ATAD1 mutation impairs postsynaptic AMPA receptor trafficking and causes a lethal encephalopathy. Brain 141, 651–661 (2018).

18. M. L. Wohlever, A. Mateja, P. T. McGilvray, K. J. Day, R. J. Keenan, Msp1 Is a Membrane Protein Dislocase for Tail-Anchored Proteins. Mol Cell 67, 194–202.e196 (2017).

19. L. Wang, H. Toutkoushian, V. Belyy, C. Y. Kokontis, P. Walter, Conserved structural elements specialize ATAD1 as a membrane protein extraction machine. Elife 11, (2022).

20. M. Babst, B. Wendland, E. J. Estepa, S. D. Emr, The Vps4p AAA ATPase regulates membrane association of a Vps protein complex required for normal endosome function. EMBO J 17, 2982–2993 (1998).

21. P. D. George Kwabena Essien Umanah et al. (Cell Reports (in press), September 2020).

22. M. L. Bochman, A. Schwacha, The Mcm complex: unwinding the mechanism of a replicative helicase. Microbiol Mol Biol Rev 73, 652–683 (2009).

23. Y. Zhai et al., Unique Roles of the Non-identical MCM Subunits in DNA Replication Licensing. Mol Cell 67, 168–179 (2017).

24. G. Cannone, S. Visentin, A. Palud, G. Henneke, L. Spagnolo, Structure of an octameric form of the minichromosome maintenance protein from the archaeon Pyrococcus abyssi. Sci Rep 7, 42019 (2017).

25. F. E. Demircioglu et al., The AAA + ATPase TorsinA polymerizes into hollow helical tubes with 8.5 subunits per turn. Nat Commun 10, 3262 (2019).

26. M. H. Ali, B. Imperiali, Protein oligomerization: how and why. Bioorg Med Chem 13, 5013–5020 (2005).

27. D. S. Goodsell, A. J. Olson, Structural symmetry and protein function. Annu Rev Biophys Biomol Struct 29, 105–153 (2000).

28. B. Tomkinson, Tripeptidyl-peptidase II: Update on an oldie that still counts. Biochimie 166, 27–37 (2019).

29. A. Fedorov et al., NCI Imaging Data Commons. Cancer Res 81, 4188–4193 (2021).

30. A. Punjani, J. L. Rubinstein, D. J. Fleet, M. A. Brubaker, cryoSPARC: algorithms for rapid unsupervised cryo-EM structure determination. Nat Methods 14, 290–296 (2017).

31. S. H. Scheres, RELION: implementation of a Bayesian approach to cryo-EM structure determination. J Struct Biol 180, 519–530 (2012).

32. E. Palovcak, D. Asarnow, M. G. Campbell, Z. Yu, Y. Cheng, Enhancing the signal-to-noise ratio and generating contrast for cryo-EM images with convolutional neural networks. IUCrJ 7, 1142–1150 (2020).

33. R. Sanchez-Garcia et al., DeepEMhancer: a deep learning solution for cryo-EM volume post-processing. Commun Biol 4, 874 (2021).

34. E. F. Pettersen et al., UCSF Chimera--a visualization system for exploratory research and analysis. J Comput Chem 25, 1605–1612 (2004).

35. T. I. Croll, ISOLDE: a physically realistic environment for model building into low-resolution electron-density maps. Acta Crystallogr D Struct Biol 74, 519–530 (2018).

36. E. F. Pettersen et al., UCSF ChimeraX: Structure visualization for researchers, educators, and developers. Protein Sci 30, 70–82 (2021).

37. T. D. Goddard et al., UCSF ChimeraX: Meeting modern challenges in visualization and analysis. Protein Sci 27, 14–25 (2018).

38. P. Emsley, B. Lohkamp, W. G. Scott, K. Cowtan, Features and development of Coot. Acta Crystallogr D Biol Crystallogr 66, 486–501 (2010).

39. P. D. Adams et al., PHENIX: a comprehensive Python-based system for macromolecular structure solution. Acta Crystallogr D Biol Crystallogr 66, 213–221 (2010).

40. V. B. Chen et al., MolProbity: all-atom structure validation for macromolecular crystallography. Acta Crystallogr D Biol Crystallogr 66, 12–21 (2010).

41. A. R. Tee, B. D. Manning, P. P. Roux, L. C. Cantley, J. Blenis, Tuberous sclerosis complex gene products, Tuberin and Hamartin, control mTOR signaling by acting as a GTPase-activating protein complex toward Rheb. Curr Biol 13, 1259–1268 (2003).

